# MrParse: Finding homologues in the PDB and the EBI AlphaFold database for Molecular Replacement and more

**DOI:** 10.1101/2021.09.02.458604

**Authors:** Adam J. Simpkin, Jens M. H. Thomas, Ronan M. Keegan, Daniel J. Rigden

## Abstract

Crystallographers have an array of search model options for structure solution by Molecular Replacement (MR). Well-established options of homologous experimental structures and regular secondary structure elements or motifs are increasingly supplemented by computational modelling. Such modelling may be carried out locally or use pre-calculated predictions retrieved from databases such as the EBI AlphaFold database. MrParse is a new pipeline to help streamline the decision process in MR by consolidating bioinformatic predictions in one place. When reflection data are provided, MrParse can rank any homologues found using eLLG which indicates the likelihood that a given search model will work in MR. In-built displays of predicted secondary structure, coiled-coil and transmembrane regions further inform the choice of MR protocol. MrParse can also identify and rank homologues in the EBI AlphaFold database, a function that will also interest other structural biologists and bioinformaticians.

## 1. Introduction

The dominant approach to solving the phase problem in crystallography is Molecular Replacement (MR). At the time of writing, 86% of crystal structures deposited in the Protein Data Bank (PDB) (Burley *et al*., 2021) in 2021 were solved by this method. By MR, initial phase estimates are derived from the placement of a search model in the asymmetric unit, typically by successive rotation and translation steps (Scapin, 2013). Successful placement requires that the search model bear a sufficiently close structural resemblance to (part of) the target structure. Conventional MR typically deploys experimental PDB structures that are inferred to be homologous to the target structure (or one of its chains or domains). The inference of homology, from a significant result in a sequence-based database search with the target as query, allows for a reasonable supposition of structural similarity of the target and PDB deposit although that assumption can break down where a protein family can adopt distinct conformations. Furthermore, with distant homologues the degree of structural similarity between target and search model may be too low for successful placement, even with advanced Maximum Likelihood-based methods (McCoy, 2004; McCoy *et al*., 2007; Read, 2001) and the use of methods to maximise their value (Rigden *et al*., 2018; Sammito *et al*., 2014).

Unconventional MR generally uses bioinformatics predictions to suggest or construct search models. Thus, a detailed consideration of the sequence properties of the target can help direct structure solution strategy (Pereira & Alva, 2021). For example, a secondary structure prediction can point to simple regular structural elements such as α-helices (Rodríguez *et al*., 2012) or recurring tertiary packing features composed of several such elements (Sammito *et al*., 2013) as potential search models. Novel and divergent folds can also be explicitly predicted using *ab initio* modelling (also known as *de novo*, free or template-independent modelling). The first broadly successful algorithms in the field (Leaver-Fay *et al*., 2011; Xu & Zhang, 2012) used fragment assembly approaches, limiting their application to relatively small targets. Limited accuracy also meant their results often needed sampling across a range of ensembles and rational edits in order to succeed in MR (Rigden *et al*., 2008; Bibby *et al*., 2012). However, *ab initio* modelling methods have advanced with remarkable speed, first by exploiting the residue contact information available from sequence alignments eg (Marks *et al*., 2011) and then, dramatically, using bespoke deep neural networks (Senior *et al*., 2020; Jumper *et al*., 2021). CASP14 saw the stunning performance of AlphaFold 2 (AF2) which, in many cases, produced predictions that resembled the target as closely as a different crystal form typically would (Pereira *et al*., 2021). The value of predictions from AF2 and the AF2-inspired RoseTTAFold (Baek *et al*., 2021) as search models was quickly demonstrated although some cases still required domain splitting or other editing (Millán *et al*., 2021; Baek *et al*., 2021; McCoy *et al*., 2021; Pereira *et al*., 2021).

As *ab initio* modelling methods have advanced, so have corresponding databases of structure predictions. Earlier efforts typically sampled uncharacterised fold space using Pfam domain definitions (Mistry *et al*., 2021) as a convenient foundation (Ovchinnikov *et al*., 2017; Lamb *et al*., 2019; Wang *et al*., 2019). Although Pfam domain boundaries inferred from sequence alignment alone are not always accurately defined, the entries in these databases could, especially with ensembling, succeed as search models (Simpkin *et al*., 2019). More recently AF2 has been used to model complete sequences of 21 whole proteomes, including human (Tunyasuvunakool *et al*., 2021) and the results made available in the EBI AlphaFold database (AFDB) (https://alphafold.ebi.ac.uk/). The often high accuracy of the predictions (and they are accompanied by high-quality residue-level error estimates) makes the database a very significant new source of search models for MR.

Here we present MrParse which addresses a number of issues in MR. It will find and rank search models from both the PDB and the AFDB providing convenient visualization of the results. It also guides choices in unconventional MR through a secondary structure prediction and predictions of regions that are relevant to MR strategy such as coiled-coil (Thomas *et al*., 2015, 2020; Caballero *et al*., 2018) and transmembrane helices. When MrParse is provided with diffraction data information it can flag the crystal pathologies that can hinder successful MR (Sevvana *et al*., 2019; Caballero *et al*., 2021) and rank experimental homologues from the PDB according to eLLG (Oeffner *et al*., 2018), a good predictor of their suitability as search models.

## 2. Methods

### 2.1. Reflection data classification

If a reflection file is provided, MrParse creates a table providing information from the reflection file (resolution, space group) and information about the crystal pathology calculated with CTRUNCATE (Evans, 2011) (non-crystallographic symmetry, twinning and anisotropy).

### 2.2. PDB database search

MrParse uses phmmer (Eddy, 2011) to search either a non-redundant or a 5% redundancy removed derivative of the PDB provided by MrBUMP (Keegan *et al*., 2018). Phmmer also provides information about the regions in the target protein that the hits correspond to. This is used to create a visualisation of the search results using Pfam Domain Graphics (Mistry *et al*., 2021) that allows for easy interpretation of how much of the target the search model covers. If a reflection file is provided, Phaser (Oeffner *et al*., 2018) is used to calculate the eLLG for each of the hits identified by phmmer. It has been shown that eLLG is a better indicator of whether a search model will succeed in MR than sequence identity (Oeffner *et al*., 2018). Therefore, when a reflection file is provided, the search results are ranked by eLLG. Any hits are downloaded from the PDB and trimmed according to their match to the target sequence.

### 2.3. Protein classification

MrParse performs protein classification analysis on the input sequence to predict secondary structure, transmembrane regions, and coiled-coil regions. Secondary structure is predicted using the Jpred4 (Drozdetskiy *et al*., 2015) RESTful Application Programming Interface (API), transmembrane regions are predicted by TMHMM (Krogh *et al*., 2001) and coiled-coil regions are predicted by deepcoil (Ludwiczak *et al*., 2019). Currently, coiled-coil and transmembrane predictions require local installations of TMHMM and deepcoil.

### 2.4. EBI AlphaFold database search

MrParse uses phmmer to search the sequence database provided by the EBI AlphaFold database (https://alphafold.ebi.ac.uk). As in the PDB database search, information from phmmer is used to create a visualisation of the search results using Pfam Domain Graphics. For the EBI AlphaFold database, these visualisations are coloured by pLDDT on an orange to blue scale, where orange indicates very low confidence in the model and blue indicates very high confidence in the model. Additional information is provided about the quality of the AF2 models, including the average pLDDT and a new measure of structural quality called the H-score.

H-score can be calculated with the following equation, where *N* represents a list of pLDDT scores and ∥*N*∥ represents the number of elements in *N*:

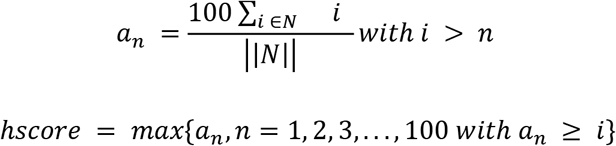

Any hits are downloaded from the database and trimmed according to their match to the target sequence and the pLDDT scores are converted into estimated B-factors using an algorithm developed for *phaser.voyager* (Claudia Milán, https://gitlab.developers.cam.ac.uk/scm/haematology/readgroup/phaser_voyager/-/blob/master/src/Voyager/MDSLibraries/pdb_structure.py). Interpreting pLDDT as B-factors improves the likelihood of success in MR by downweighting the less reliable regions of the model (Croll *et al*., 2019).

## 3. Examples

### 3.1. Interpreting MrParse report page

Figure 1 shows an example of a MrParse report page generated from the reflection data and sequence data for PDB entry 5LM4. Here, we use 5LM4 to demonstrate how to interpret the results of an MrParse run.

**Figure 1):**
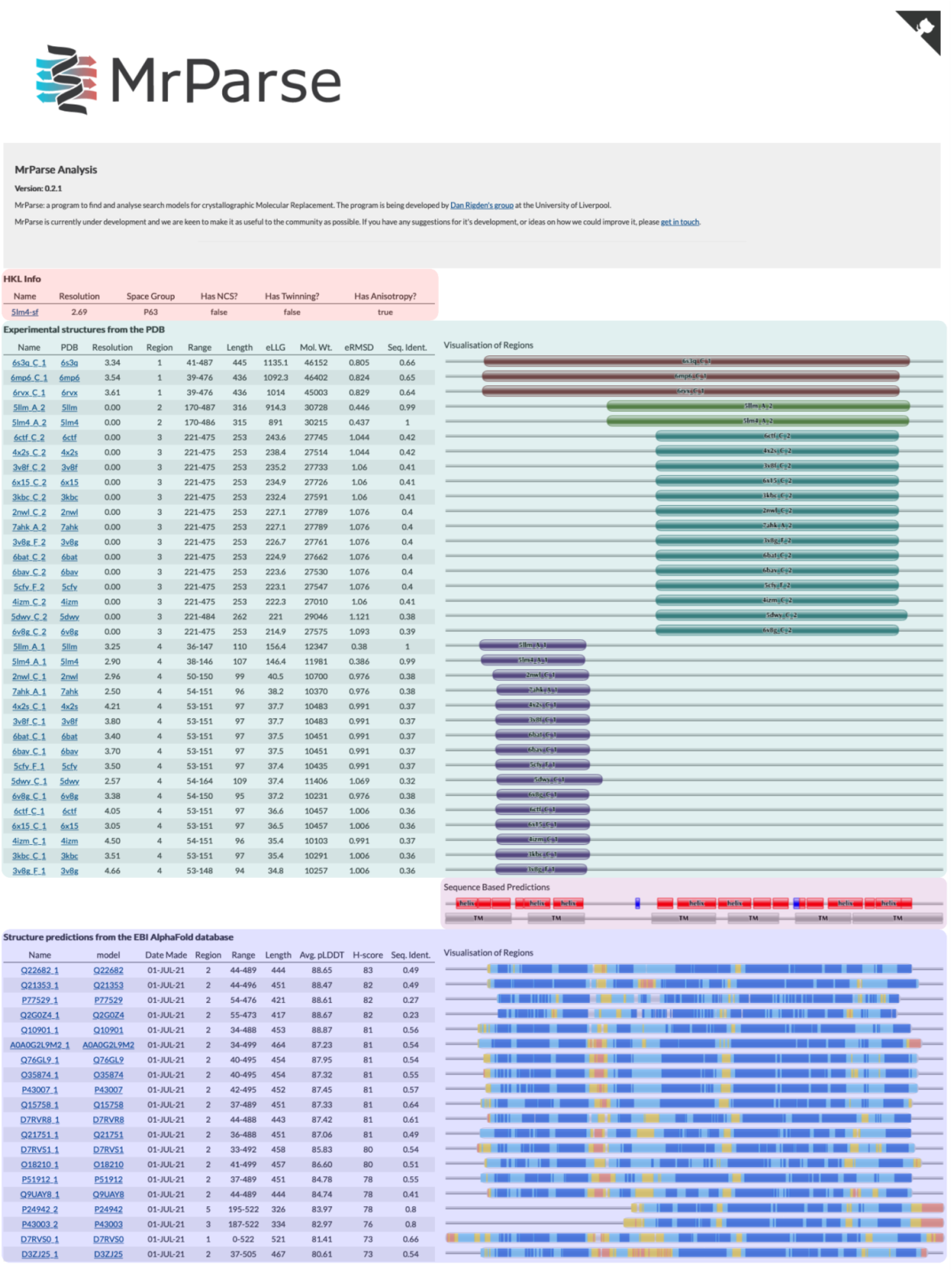
Highlighted sections of an MrParse report page. In red is information on the input reflection file, including resolution, space group and crystal pathology. In teal is information about the PDB entries identified by Phmmer and visualisations of the matches. In purple is the protein classification report, this includes a secondary structure prediction, a coiled-coil prediction, and a transmembrane prediction. Finally, in blue is information about the AlphaFold models identified by Phmmer and visualisations of the matches coloured by pLDDT on an orange to blue scale, where orange indicates very low confidence in the model and blue indicates very high confidence in the model.

#### 3.1.1 HKL info

The ‘HKL info’ panel (Figure 1, Red) allows us to assess whether there are any crystal pathologies that might make MR more difficult. For example, the detection of translational non-crystallographic symmetry can be important for successful MR (Caballero *et al*., 2021). In the case of 5LM4, we have a 2.69Å dataset which shows anisotropy. Phaser can be used to correct anisotropic data and performs this step automatically in its autoMR pipeline (McCoy *et al*., 2007).

#### 3.1.2 Experimental structures from the PDB

The ‘Experimental structures from the PDB’ panel (Figure 1, Teal) provides information about homologues identified by Phmmer. In this example we can see that we have identified three near full length matches when looking at the visualisation of regions on the right-hand side (PDB entries: 6S3Q, 6MP6, 6RVX). These hits all have high sequence identity to our target (65%, 66% and 64% respectively) and give high eLLG scores (1135.1, 1092.3 and 1014 respectively). When eLLG is much greater than 60, structure solution by MR is likely to be straightforward (Oeffner *et al*., 2018), therefore we can be fairly confident that these search models will work in MR. Further down the list of hits it can be seen that the target seems to match experimental structures in two distinct regions, likely corresponding to structural domains. Any matches are downloaded from the PDB and trimmed to match the target sequence. These are downloaded into the ‘homologues’ subdirectory in the MrParse run directory.

#### 3.1.3 Sequence based predictions

The ‘Sequence based predictions’ panel (Figure 1, Purple) provides secondary structure, transmembrane and coiled-coil predictions. In this example, Jpred4 predicts a large number of helices and TMHMM predicts several transmembrane regions. For a high resolution dataset that is predicted to be predominantly helical, an approach such as AMPLE helical ensembles (Sánchez Rodríguez *et al*., 2020) or ARCIMBOLDO (Rodríguez *et al*., 2012) can be used. If coiled-coils were predicted, AMPLE and ARCIMBOLDO also have coiled-coil specific modes that can be tried (Thomas *et al*., 2020; Caballero *et al*., 2018).

#### 3.1.4 Structure predictions from the EBI AlphaFold database

The ‘Structure predictions from the EBI AlphaFold database’ panel (Figure 1, Blue) provides information about AF2 models identified by Phmmer in the AFDB. In this example we can see a large number of AF2 hits. These hits are largely very high quality with an average pLDDT score >80 for all of the hits. The visualisation on the right-hand side shows the regions that the models correspond to and provides information about predicted model reliability at a residue level. For example, the few models that match the C-terminal region of the target structure (P24942, P43003 & D7RVS0) all have lower predicted reliability in the region. Any matches are downloaded from the AFDB and trimmed to match the target sequence and undergo a pLDDT to estimated B-factor conversion to improve their performance in MR. These are downloaded into the ‘models’ subdirectory in the MrParse run directory.

### 3.2. Use of AFDB entry for MR when PDB search model is lacking

7DRY is a crystal structure of *Aspergillus oryzae* Rib2 deaminase experimentally determined by Zn-SAD (Chen *et al*., 2021). A phmmer search of the PDB only identified a single hit (PDB code: 2cvi) that only covers a 71 residue region of the target protein with 31% sequence identity (Figures 2A & 2B). This homologue was insufficiently similar to the target protein to succeed in MR. A search of the EBI AlphaFold2 database identified a number of models that covered a larger region of the target protein and with a higher sequence identity. MR with the model of Q12362, the best hit ranked by H-score (Figure 2A & 2C), was successfully placed by Phaser (LLG: 173, TFZ: 15.4) and rebuilt with Buccaneer (Cowtan, 2006) (R-factor: 0.23, R-free: 0.25).

**Figure 2).**
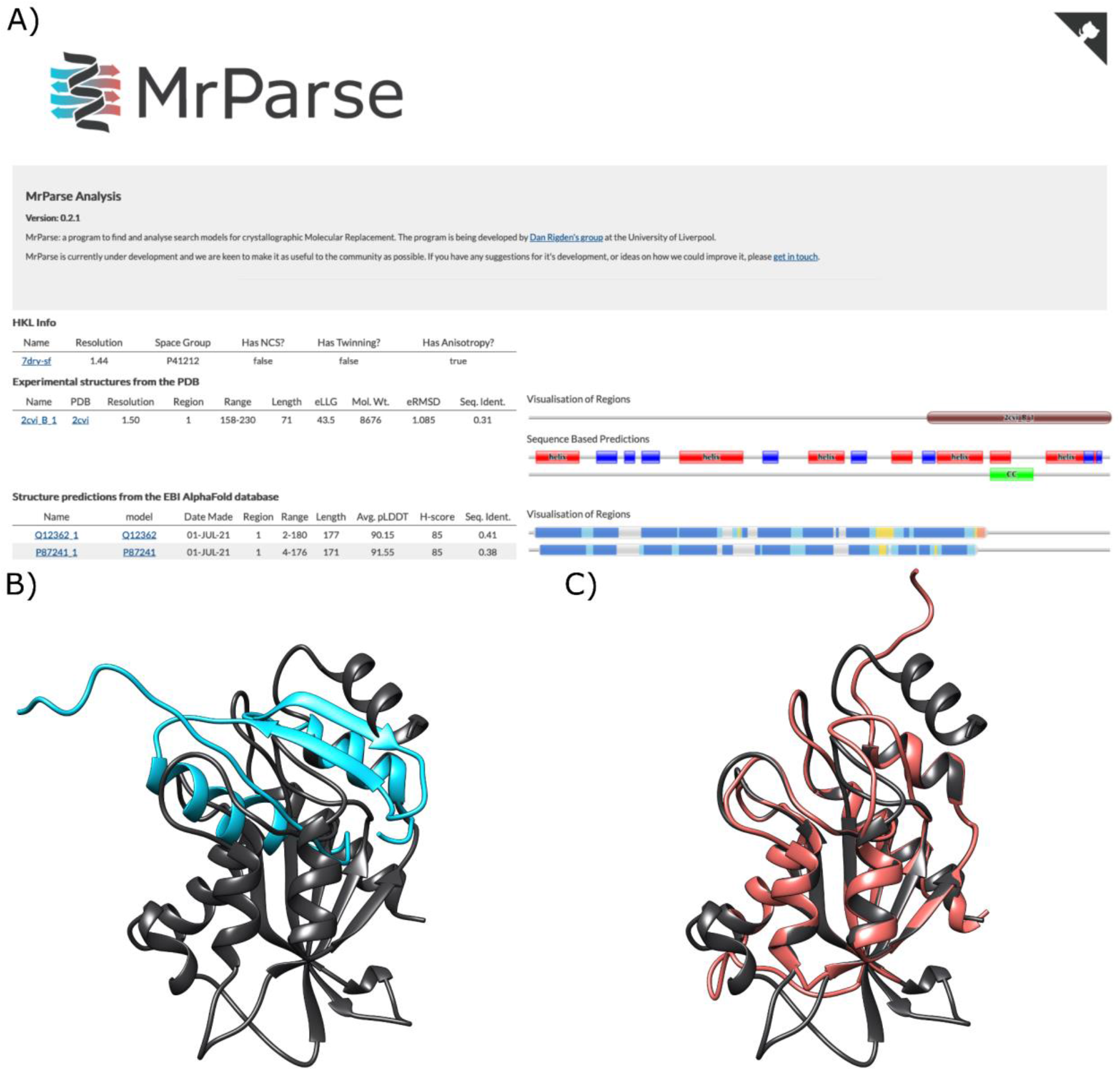
A - MrParse results page, components are as seen previously except for a coiled-coil prediction (labelled CC) under the sequence-based prediction heading. B - Closest match in the PDB (2CIV, blue) aligned to the crystal structure (7DRY, grey). C - Closest match in the EBI AlphaFold database (Q12362, coral) aligned to the crystal structure (7DRY, grey).

## 4. Discussion

A crystallographer attempting to solve a macromolecular crystal structure by MR should be aware of the existence of any crystal pathologies and has an increasing range of search model options to choose from. MrParse is designed to bring together a range of relevant information in a single place and present it with useful visualisations and sortable tables. For most effective use, it expects both diffraction data and a target sequence, but it can run without the former. Conventional MR using homologous structures identified in the PDB is supported by presentation of potential search models, discovered by Phmmer, with graphics that illustrate their extent relative to the target, and numerical data that illustrate their size and characteristics. In the future more sensitive HHpred (Soding, 2005) sequence searching will be supported. With diffraction data supplied, search models are ordered by default by eLLG as a predictor of their relative utility in MR. At present, PDB files are available locally and through the CCP4i2 GUI (Potterton *et al*., 2018): future work will integrate MrParse better into the CCP4 Cloud setting (Krissinel *et al*., 2018). In the future, options for inline composition of ensembles will be implemented. The PDB files, which are trimmed according to their match to the target sequence, and modified to convert the predicted residue error into a B-factor (Claudia Milán, https://gitlab.developers.cam.ac.uk/scm/haematology/readgroup/phaser_voyager/-/blob/master/src/Voyager/MDSLibraries/pdb_structure.py), can be fed directly to programs such as Phaser (McCoy *et al*., 2007) or MOLREP (Vagin & Teplyakov, 2010) or may, in more difficult cases, require special treatment (Vagin & Teplyakov, 2010; Rigden *et al*., 2018; Simpkin *et al*., 2019; Sammito *et al*., 2014). The also well-established use of secondary structure elements as search models (Rodríguez *et al*., 2012), especially at higher resolution, is also facilitated by secondary structure prediction that enables, for example, helpful predictions of likely helix lengths (Rodríguez *et al*., 2012).

Perhaps the most exciting and forward-facing aspect of MrParse is its discovery of structure predictions, especially those generated by *ab initio* (also known as *de novo* or template-independent) methods. The potential of these methods for MR of targets with novel or divergent folds has been recognised for some time (Rigden *et al*., 2008; Bibby *et al*., 2012; Qian *et al*., 2007). Nevertheless, their (until recently) considerable CPU demands and specialist software have undoubtedly proved off-putting to structural biologists, despite the convenience offered by some servers (Keegan *et al*., 2015). In addition, the accuracy of *ab initio* methods has, historically, not always been sufficient for MR and only smaller proteins were tractable by the earliest methods. This picture has changed rapidly in recent years with first AlphaFold (Senior *et al*., 2020) and then AlphaFold2 (Jumper *et al*., 2021), each providing a step-change in model accuracy. These developments have been mirrored in online databases of *ab initio* structure predictions. Databases based on earlier methods such as GREMLIN (Ovchinnikov *et al*., 2017), PconsFam (Lamb *et al*., 2019) and C-QUARK (Wang *et al*., 2019) typically modelled single representatives of Pfam families. These provided useful sampling of uncharacterised protein fold space, sometimes being suitable for MR (Simpkin *et al*., 2019), but were limited by the fact that the domain boundaries of Pfam entries are not always, in the absence of some kind of structural information, accurately determined from sequence analysis (Bateman *et al*., 2010). The AFDB, in contrast, includes full-length models from 21 essentially complete proteomes, with the ambition to cover UniRef90 (Suzek *et al*., 2015) - so that no protein of interest will be less than 90% identical to an entry in the database - by the end of 2021. Models in the AFDB are likely to be much more accurate than models available elsewhere and are accompanied by accurate residue-level error estimates. Their availability therefore has profound implications for the choice of crystallographic phasing method (Kryshtafovych *et al*., 2021; McCoy *et al*., 2021) and the already very high market share of MR will only increase further.

MrParse currently provides a second graphical panel devoted solely to matches in the AFDB. These can be ranked by clicking on column headings for two measures of model quality, including the novel H-score described here, or percentage sequence identity between the protein of interest and the model. While the experience of the CASP structure prediction experiment suggests that many models serve, unaltered, as successful search models, downstream editing of models after retrieval via MrParse will sometimes be necessary (McCoy *et al*., 2021; Millán *et al*., 2021; Pereira *et al*., 2021). This can eliminate regions with low predicted accuracy (McCoy *et al*., 2021) or sample a variety of truncated versions (Pereira *et al*., 2021); or to excise domains from multi-domain models, recognising that inter-domain packing remains a challenge for AF. Presently, these hits are found by phmmer (Eddy, 2011) search against a local database containing the sequences of entries in the AFDB. With the ambitious plans to expand the AF database this arrangement becomes increasingly awkward as ever-larger databases would need to be distributed with CCP4. Happily, the 3D-Beacons initiative (Orengo *et al*., 2020) will shortly be launching an API for sequence-based retrieval of models, not only from the AFDB, but also from a variety of other resources containing protein structure predictions. Thus, we envisage that the importance of MrParse in facilitating access to a wide range of potential MR search models, both experimental structures and predictions, will only grow in the future. In addition, its ability to search AFDB in particular and conveniently visualise the results is likely to prove useful to bioinformaticians and cryo-EM researchers (Kryshtafovych *et al*., 2021; Simpkin *et al*. 2021) as well as to crystallographers.

## 5 Acknowledgements

This work was supported by the Biotechnology and Biological Sciences Research Council [BB/S007105/1] and by a CCP4 grant to AJS.

## Bibliography

AlphaFold Protein Structure Database (2021). AlphaFold Protein Structure Database, https://alphafold.ebi.ac.uk

Baek, M., DiMaio, F., Anishchenko, I., Dauparas, J., Ovchinnikov, S., Lee, G. R., Wang, J., Cong, Q., Kinch, L. N., Schaeffer, R. D., Millán, C., Park, H., Adams, C., Glassman, C. R., DeGiovanni, A., Pereira, J. H., Rodrigues, A. V., van Dijk, A. A., Ebrecht, A. C., Opperman, D. J., Sagmeister, T., Buhlheller, C., Pavkov-Keller, T., Rathinaswamy, M. K., Dalwadi, U., Yip, C. K., Burke, J. E., Garcia, K. C., Grishin, N. V., Adams, P. D., Read, R. J. & Baker, D. (2021). Science.

Bateman, A., Coggill, P. & Finn, R. D. (2010). Acta Crystallogr. Sect. F Struct. Biol. Cryst. Commun. 66, 1148–1152.

Bibby, J., Keegan, R. M., Mayans, O., Winn, M. D. & Rigden, D. J. (2012). Acta Crystallogr. D Biol. Crystallogr. 68, 1622–1631.

Burley, S. K., Bhikadiya, C., Bi, C., Bittrich, S., Chen, L., Crichlow, G. V., Christie, C. H., Dalenberg, K., Di Costanzo, L., Duarte, J. M., Dutta, S., Feng, Z., Ganesan, S., Goodsell, D. S., Ghosh, S., Green, R. K., Guranović, V., Guzenko, D., Hudson, B. P., Lawson, C. L., Liang, Y., Lowe, R., Namkoong, H., Peisach, E., Persikova, I., Randle, C., Rose, A., Rose, Y., Sali, A., Segura, J., Sekharan, M., Shao, C., Tao, Y.-P., Voigt, M., Westbrook, J. D., Young, J. Y., Zardecki, C. & Zhuravleva, M. (2021). Nucleic Acids Res. 49, D437–D451.

Caballero, I., Sammito, M. D., Afonine, P. V., Usón, I., Read, R. J. & McCoy, A. J. (2021). Acta Crystallogr D Struct Biol. 77, 131–141.

Caballero, I., Sammito, M., Millán, C., Lebedev, A., Soler, N. & Usón, I. (2018). Acta Crystallogr D Struct Biol. 74, 194–204.

Chen, S.-C., Ye, L.-C., Yen, T.-M., Zhu, R.-X., Li, C.-Y., Chang, S.-C., Liaw, S.-H. & Hsu, C.-H. (2021). IUCrJ. 8, 549–558.

Cowtan, K. (2006). Acta Crystallogr. D Biol. Crystallogr. 62, 1002–1011.

Croll, T. I., Sammito, M. D., Kryshtafovych, A. & Read, R. J. (2019). Proteins. 87, 1113–1127.

Drozdetskiy, A., Cole, C., Procter, J. & Barton, G. J. (2015). Nucleic Acids Res. 43, W389–W394.

Eddy, S. R. (2011). PLoS Comput. Biol. 7, e1002195.

Evans, P. R. (2011). Acta Crystallogr. D Biol. Crystallogr. 67, 282–292.

Jumper, J., Evans, R., Pritzel, A., Green, T., Figurnov, M., Ronneberger, O., Tunyasuvunakool, K., Bates, R., Žídek, A., Potapenko, A., Bridgland, A., Meyer, C., Kohl, S. A. A., Ballard, A. J., Cowie, A., Romera-Paredes, B., Nikolov, S., Jain, R., Adler, J., Back, T., Petersen, S., Reiman, D., Clancy, E., Zielinski, M., Steinegger, M., Pacholska, M., Berghammer, T., Bodenstein, S., Silver, D., Vinyals, O., Senior, A. W., Kavukcuoglu, K., Kohli, P. & Hassabis, D. (2021). Nature.

Keegan, R. M., Bibby, J., Thomas, J., Xu, D., Zhang, Y., Mayans, O., Winn, M. D. & Rigden, D. J. (2015). Acta Crystallogr. D Biol. Crystallogr. 71, 338–343.

Keegan, R. M., McNicholas, S. J., Thomas, J. M. H., Simpkin, A. J., Simkovic, F., Uski, V., Ballard, C. C., Winn, M. D., Wilson, K. S. & Rigden, D. J. (2018). Acta Crystallogr D Struct Biol. 74, 167–182.

Krissinel, E., Lebedev, A., Ballard, C., Uski, V. & Keegan, R. (2018). Acta Crystallographica Section A Foundations and Advances. 74, e411–e412.

Krogh, A., Larsson, B., von Heijne, G. & Sonnhammer, E. L. (2001). J. Mol. Biol. 305, 567–580.

Kryshtafovych, A., Moult, J., Albrecht, R., Chang, G. A., Chao, K., Fraser, A., Greenfield, J., Hartmann, M. D., Herzberg, O., Josts, I., Leiman, P. G., Linden, S. B., Lupas, A. N., Nelson, D. C., Rees, S. D., Shang, X., Sokolova, M. L., Tidow, H. & Team, A. F. (2021). Proteins.

Lamb, J., Jarmolinska, A. I., Michel, M., Menéndez-Hurtado, D., Sulkowska, J. I. & Elofsson, A. (2019). J. Mol. Biol. 431, 2442–2448.

Leaver-Fay, A., Tyka, M., Lewis, S. M., Lange, O. F., Thompson, J., Jacak, R., Kaufman, K., Renfrew, P. D., Smith, C. A., Sheffler, W., Davis, I. W., Cooper, S., Treuille, A., Mandell, D. J., Richter, F., Ban, Y.-E. A., Fleishman, S. J., Corn, J. E., Kim, D. E., Lyskov, S., Berrondo, M., Mentzer, S., Popović, Z., Havranek, J. J., Karanicolas, J., Das, R., Meiler, J., Kortemme, T., Gray, J. J., Kuhlman, B., Baker, D. & Bradley, P. (2011). Methods Enzymol. 487, 545–574.

Ludwiczak, J., Winski, A., Szczepaniak, K., Alva, V. & Dunin-Horkawicz, S. (2019). Bioinformatics. 35, 2790–2795.

Marks, D. S., Colwell, L. J., Sheridan, R., Hopf, T. A., Pagnani, A., Zecchina, R. & Sander, C. (2011). PLoS One. 6, e28766.

McCoy, A. J. (2004). Acta Crystallographica Section D Biological Crystallography. 60, 2169–2183.

McCoy, A. J., Grosse-Kunstleve, R. W., Adams, P. D., Winn, M. D., Storoni, L. C. & Read, R. J. (2007). J. Appl. Crystallogr. 40, 658–674.

McCoy, A. J., Sammito, M. D. & Read, R. J. (2021). Possible Implications of AlphaFold2 for Crystallographic Phasing by Molecular Replacement.

Millán, C., Keegan, R. M., Pereira, J., Sammito, M. D., Simpkin, A. J., McCoy, A. J., Lupas, A. N., Hartmann, M. D., Rigden, D. J. & Read, R. J. (2021). Proteins.

Millán, C. (2021). Phaser Voyager GitLab, https://gitlab.developers.cam.ac.uk/scm/haematology/readgroup/phaser_voyager/-/blob/master/src/Voyager/MDSLibraries/pdb_structure.py

Mistry, J., Chuguransky, S., Williams, L., Qureshi, M., Salazar, G. A., Sonnhammer, E. L. L., Tosatto, S. C. E., Paladin, L., Raj, S., Richardson, L. J., Finn, R. D. & Bateman, A. (2021). Nucleic Acids Res. 49, D412–D419.

Oeffner, R. D., Afonine, P. V., Millán, C., Sammito, M., Usón, I., Read, R. J. & McCoy, A. J. (2018). Acta Crystallographica Section D: Structural Biology. 74, 245–255.

Orengo, C., Velankar, S., Wodak, S., Zoete, V., Bonvin, A. M. J. J., Elofsson, A., Feenstra, K. A., Gerloff, D. L., Hamelryck, T., Hancock, J. M., Helmer-Citterich, M., Hospital, A., Orozco, M., Perrakis, A., Rarey, M., Soares, C., Sussman, J. L., Thornton, J. M., Tuffery, P., Tusnady, G., Wierenga, R., Salminen, T. & Schneider, B. (2020). F1000Res. 9,.

Ovchinnikov, S., Park, H., Varghese, N., Huang, P.-S., Pavlopoulos, G. A., Kim, D. E., Kamisetty, H., Kyrpides, N. C. & Baker, D. (2017). Science. 355, 294–298.

Pereira, J. & Alva, V. (2021). Acta Crystallographica Section D Structural Biology. 77,.

Pereira, J., Simpkin, A. J., Hartmann, M. D., Rigden, D. J., Keegan, R. M. & Lupas, A. N. (2021). Proteins: Structure, Function, and Bioinformatics.

Potterton, L., Agirre, J., Ballard, C., Cowtan, K., Dodson, E., Evans, P. R., Jenkins, H. T., Keegan, R., Krissinel, E., Stevenson, K., Lebedev, A., McNicholas, S. J., Nicholls, R. A., Noble, M., Pannu, N. S., Roth, C., Sheldrick, G., Skubak, P., Turkenburg, J., Uski, V., von Delft, F., Waterman, D., Wilson, K., Winn, M. & Wojdyr, M. (2018). Acta Crystallogr D Struct Biol. 74, 68–84.

Qian, B., Raman, S., Das, R., Bradley, P., McCoy, A. J., Read, R. J. & Baker, D. (2007). Nature. 450, 259–264.

Read, R. J. (2001). Acta Crystallogr. D Biol. Crystallogr. 57, 1373–1382.

Rigden, D. J., Keegan, R. M. & Winn, M. D. (2008). Acta Crystallogr. D Biol. Crystallogr. 64, 1288–1291.

Rigden, D. J., Thomas, J. M. H., Simkovic, F., Simpkin, A., Winn, M. D., Mayans, O. & Keegan, R. M. (2018). Acta Crystallogr D Struct Biol. 74, 183–193.

Rodríguez, D., Sammito, M., Meindl, K., de Ilarduya, I. M., Potratz, M., Sheldrick, G. M. & Usón, I. (2012). Acta Crystallographica Section D Biological Crystallography. 68, 336–343.

Sammito, M., Meindl, K., de Ilarduya, I. M., Millán, C., Artola-Recolons, C., Hermoso, J. A. & Usón, I. (2014). FEBS J. 281, 4029–4045.

Sammito, M., Millán, C., Rodríguez, D. D., de Ilarduya, I. M., Meindl, K., De Marino, I., Petrillo, G., Buey, R. M., de Pereda, J. M., Zeth, K., Sheldrick, G. M. & Usón, I. (2013). Nature Methods. 10, 1099–1101.

Sánchez Rodríguez, F., Simpkin, A. J., Davies, O. R., Keegan, R. M. & Rigden, D. J. (2020). Acta Crystallographica Section D: Structural Biology. 76, 962–970.

Scapin, G. (2013). Acta Crystallogr. D Biol. Crystallogr. 69, 2266–2275.

Senior, A. W., Evans, R., Jumper, J., Kirkpatrick, J., Sifre, L., Green, T., Qin, C., Žídek, A., Nelson, A. W. R., Bridgland, A., Penedones, H., Petersen, S., Simonyan, K., Crossan, S., Kohli, P., Jones, D. T., Silver, D., Kavukcuoglu, K. & Hassabis, D. (2020). Nature. 577, 706–710.

Sevvana, M., Ruf, M., Usón, I., Sheldrick, G. M. & Herbst-Irmer, R. (2019). Acta Crystallogr D Struct Biol. 75, 1040–1050.

Simpkin, A. J., Thomas, J. M. H., Simkovic, F., Keegan, R. M. & Rigden, D. J. (2019). Acta Crystallogr D Struct Biol. 75, 1051–1062.

Simpkin, A. J., Winn, M. D., Rigden, D. J. & Keegan, R. M. (2021). Acta Crystallographica Section D: Structural Biology. In revision.

Soding, J. (2005). Bioinformatics. 21, 951–960.

Suzek, B. E., Wang, Y., Huang, H., McGarvey, P. B., Wu, C. H. & UniProt Consortium (2015). Bioinformatics. 31, 926–932.

Thomas, J. M. H., Keegan, R. M., Bibby, J., Winn, M. D., Mayans, O. & Rigden, D. J. (2015). IUCrJ. 2, 198–206.

Thomas, J. M. H., Keegan, R. M., Rigden, D. J. & Davies, O. R. (2020). Acta Crystallogr D Struct Biol. 76, 272–284.

Tunyasuvunakool, K., Adler, J., Wu, Z., Green, T., Zielinski, M., Žídek, A., Bridgland, A., Cowie, A., Meyer, C., Laydon, A., Velankar, S., Kleywegt, G. J., Bateman, A., Evans, R., Pritzel, A., Figurnov, M., Ronneberger, O., Bates, R., Kohl, S. A. A., Potapenko, A., Ballard, A. J., Romera-Paredes, B., Nikolov, S., Jain, R., Clancy, E., Reiman, D., Petersen, S., Senior, A. W., Kavukcuoglu, K., Birney, E., Kohli, P., Jumper, J. & Hassabis, D. (2021). Nature. 596, 590–596.

Vagin, A. & Teplyakov, A. (2010). Acta Crystallogr. D Biol. Crystallogr. 66, 22–25.

Wang, Y., Shi, Q., Yang, P., Zhang, C., Mortuza, S. M., Xue, Z., Ning, K. & Zhang, Y. (2019). Genome Biol. 20, 229.

Xu, D. & Zhang, Y. (2012). Proteins. 80, 1715–1735.

